# A Microfluidic Platform for Whole-Membrane Integrity Profiling in Live Neuronal Cells

**DOI:** 10.1101/2025.08.02.668278

**Authors:** Till Ryser, Ata Krichene, Nicolò Marchi, Felix Rodriguez Espinal, Anne-Laure Mahul-Mellier, Hilal A. Lashuel, Carlotta Guiducci

## Abstract

Structural and functional compromise of the cellular membrane are central mechanisms in the pathogenesis of numerous diseases, including neurodegenerative disorders such as Alzheimer’s and Parkinson’s disease. However, existing techniques for assessing membrane integrity often lack the ability to provide dynamic, whole-cell measurements and are limited to localized damage detection or population-level analysis. There is a growing need for methods capable of monitoring membrane integrity over time at the single-cell level and across the entire membrane surface. In this study, we present a microfluidic platform for real-time, label-free assessment of membrane integrity by analyzing dielectric properties. We apply this system to investigate how different aggregated forms of α-Synuclein (aSyn), a protein that plays a central role in the pathogenesis of Parkinson’s disease and disrupts neuronal membranes. Our platform integrates electrokinetic microdevices with 3D microelectrodes and imaging, enabling continuous analysis of up to 30 live neuronal cells per hour under flow. By measuring electrorotation responses, we quantify changes in plasma membrane capacitance in response to monomeric, oligomeric, and fibrillar aSyn. This approach allows direct, time-resolved comparison of membrane-disruptive effects across different aSyn conformations with single-cell resolution and whole-membrane sensitivity.

## Introduction

Membrane disruption stands out as a central, mechanism across many disorders, including muscular dystrophy^1^, multiple sclerosis^2^, and certain neurodegenerative diseases^3^. Cellular membranes are critical structures that regulate molecular transport^4^, signaling^5^, and overall cellular homeostasis^6^. In many disorders, pathological processes lead to compromised membrane integrity, either through altered lipid composition^7^, integration of toxic proteins into the membrane and pore formation^8^, or imbalanced ion gradients^9^. Recognizing membrane disruption as a common denominator underscores the need for techniques studying relevant properties of the cellular membrane, such as their physical integrity, dielectric properties, permeability, and rigidity.

Currently, dye-based approaches such as cytotoxicity assays, as well as lipid membrane systems with fluorescent dye release assays, are mostly used to study membrane damage^10^. However, dyes only indirectly measure membrane damage as they depend on the increased permeability of the membrane or changes in metabolic activity. Additionally, their two-dimensional optical readout makes it challenging to assess the extent of the damage. Techniques that directly measure the physical phenotype of individual cells offer many advantages. Atomic force microscopy (AFM) and scanning electron microscopy (SEM), on the other hand, are often used to localize nanoscale to microscale features in the cell membrane but are restricted to studying the outer surface and have low throughput, or in the case of the SEM, suffer from their limited applicability to live cells.

Electrophysiology measurements are commonly used to assess membrane properties and permeabilization, as well as membrane-mediated toxicity^11,12^. However, among these, traditional single-cell methods such as patch clamp techniques^13^ are complex and time-consuming, making them unsuitable for large-scale studies. Electrical impedance spectroscopy (EIS) and electrokinetic techniques are more promising in this regard. They are label-free and non-disruptive techniques capable of measuring biological particles in real time. Furthermore, they can be integrated into microfluidic devices to mitigate some of the challenges of throughput, sample preparation, and electrode proximity^14^. EIS measures perturbations in the local electrical environment caused by cells when mild electric fields are applied, enabling direct assessment of their dielectric properties, such as overall permittivity and membrane insulating capacity. However, to minimize parasitic effects, cells need to be positioned in close proximity to the electrodes, which often causes shear stress or cellular damage^15^. Furthermore, EIS often measures ensembles of cells simultaneously^16^. While insightful, these approaches do not provide the necessary granularity, and the variability in the measurement conditions of the single cells in the batch limits their performance. Electrokinetics, on the other hand, is an electrophysiological technique which focuses on the movement of cells in response to non-homogenous electric fields, providing insights into the dielectric and electric properties of cells. It studies the induced movement caused by polarization (dielectrophoresis) or charge (electrophoresis) of single cells by observing their behavior in a non-homogeneous electric field. The electrical fingerprint of a cell can provide valuable insights into membrane parameters such as integrity, capacitance, or conductance of the phospholipid bilayer of the plasma membrane. Additionally, the electrokinetic force can be used to position or sort the cell post measurement, offering an advantage over conventional EIS.

We propose a method combining microfluidics and electrorotation to study membrane damage at the single-cell level. In microfluidic systems, electrorotation is increasingly used for single-cell analysis^17^, distinguishing healthy from diseased cells^18^, and probing biophysical changes in response to external stimuli^19^—making it a valuable tool for diagnostics, cell sorting, and basic research. By integrating the electrokinetics principle of electrorotation into microfluidics with micropillar electrode technology, we have developed a unique system for precise electrophysiology measurements on live single cells exposed to amyloidogenic proteins. This integration not only facilitates single-cell analysis but also minimizes issues related to parasitic effects and measurement variability. Moreover, the use of microcages to immobilize cells in flow via dielectrophoresis^20^, leveraging the same cell polarization mechanism used in electrorotation, eliminates the need for optical tweezers or physical trapping methods. As the electric field varies, the cell experiences a torque dependent on its polarization and membrane integrity, leading to rotation. The resulting rotation speed can be fitted using the Clausius-Mossotti factor^21^, which allows extraction of key parameters correlated with membrane integrity, such as the membrane’s electrical capacitance.

To demonstrate the capability of our system to evaluate membrane damage, we studied and compared the membrane-disrupting properties of monomeric and aggregated α-Synuclein (aSyn), with the latter being strongly associated with neurotoxic effects. aSyn is a 14 kDa presynaptic protein whose exact physiological function remains unclear, but it has been shown to be involved in regulating neurotransmitter release and synaptic vesicle trafficking^22^. In neurodegenerative diseases, such as Parkinson’s disease (PD), Alzheimer’s disease (AD) dementia with Lewy bodies (DLB), and multiple system atrophy (MSA), it becomes misfolded and aggregates into oligomers, fibrils that accumulate and associate with other proteins, lipids, and membranous organelles to ultimately form inclusions known as Lewy bodies, which represent a pathological hallmark of synucleinopathies^23^. These aggregate species are recognized as key contributors to the pathogenesis of neurodegenerative diseases through different mechanisms, including the interaction with and disruption of cellular membrane structure and/or function. Leveraging our expertise, we use distinct, highly pure, and well-characterized forms of aSyn species (monomers, oligomers, or fibrils) to directly compare their different membrane-compromising effects using our approach.

While for the monomeric form, no or low membrane toxicity is reported^10^, the oligomeric form has been recognized as toxic by a multitude of studies^24,25^. The β-sheet-rich structure of aSyn oligomers is thought to be linked to the ability of oligomers to integrate into or disrupt the lipid bilayer of the membrane^26,27^. Oligomeric aSyn can adopt a pore structure, causing the internal interruption of ion homeostasis, leading to cell death^28,29^. Others have highlighted the key role of oligomers and their detergent-like interaction with the lipid bilayer, causing its disruption^33^.

Cell interaction with aSyn fibrils has been closely linked to neurotoxicity. The insertion of fibrils into the membrane can compromise its integrity and stability due to their interaction with the phospholipid bilayer^34^. Several studies have reported fibril-induced toxicity through reductions in cellular metabolic activity^35,36^. Interestingly, Mahul-Mellier *et al*. showed that while purified fibrils alone appeared non-toxic, their toxicity increased significantly upon the addition of excess monomeric aSyn^10^. This enhanced toxicity was attributed to fibril elongation at the membrane, where monomers act as seeds for further aggregation^10^, as well as monomer-driven secondary nucleation processes that release toxic oligomeric forms^37^.

To directly probe the impact of these different aSyn species on neuronal membranes, we developed a tri-dimensional microelectrode-based electrorotation platform capable of assessing membrane integrity at the single-cell level. Our microfluidic system supports high-throughput measurements of up to 30 live neuronal cells per hour under flow conditions. A Galton array was incorporated to enrich single cells within the measurement region and filter out larger cell aggregates. By capturing electrorotation profiles, we quantified changes in plasma membrane electrical capacitance following exposure to monomeric, oligomeric, fibrillar aSyn, and mixtures of monomers and fibrils at various time points. This single-cell approach provides a sensitive readout of membrane disruption, underscoring electrorotation as a powerful technique to investigate the membrane-damaging effects of different aggregate aSyn species.

## Results

### Single-cell electrorotation in microfluidics to assess membrane damage

Designing an electrorotation setup that achieves sufficient throughput while maintaining cell viability is critical for obtaining statistically significant measurements of cell populations. The core of the system is an array of five measurement sites where the electrokinetic behavior of single cells is simultaneously characterized.

To separate single cells from large clusters, we implemented a Galton array at the center of the channel (Fig. 1abc). Cells must be small enough to pass between the pillars of the array. Once they enter, they are stochastically guided to the center of the chip. The Galton array effectively prevents contamination and clogging in the cage region, enabling the measurement of a larger number of cells per session. This setup significantly improves throughput and reduces measurement time compared to our previously described electrorotation setup. At the end of the Galton array, five microcages, each consisting of tri-dimensional platinum electrodes (80 µm in diameter and spanning the entire 50 µm height of the channel), are used to perform electrorotation measurements of five single cells simultaneously (Fig. 1c).

**Fig. 1.**
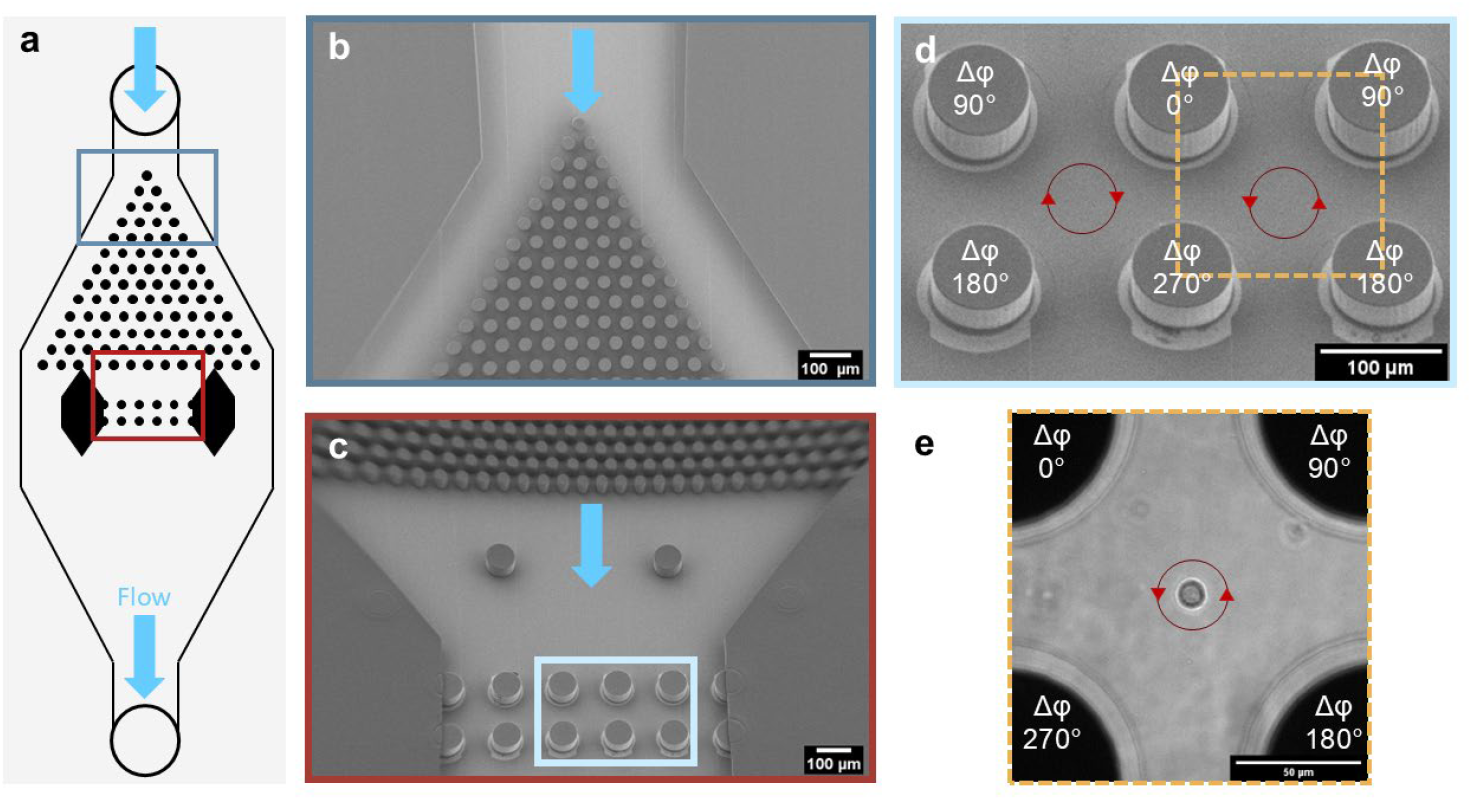
Overview of the electrorotation device. **a** Schematic of the microfluidic electrorotation chip with two highlighted sections. **b** Entrance to the Galton array in the microfluidic channel. Flow is indicated by a light blue arrow. SEM picture of the microcages and the end of the Galton array. The scale bar equals 100 µm. **c** Five microcages are used for the simultaneous recording of spectra. SU8-channels confine the space and direct the cells further toward the traps. The scale bar equals 100 µm. **d** Two adjacent electrorotation cages enable the recording of multiple cells at once. The yellow dotted square indicates an electrorotation microcage. Red arrows describe the rotation sense of the rotated cells. Each electrode applies the same AC electric field with a respective phase shift Δφ of 90 degrees The scale bar equals 100 µm. **e** Image of a live M17 neuroblastomal cell captured and rotated in a microcage. Red arrows describe the rotation sense of the cell at the frequency range applied in this experiment. Each electrode applies the same AC electric field with a respective phase shift Δφ of 90 degrees. The scale bar equals 50 µm.

When a single cell reaches a microcage, the flow is halted, and a dielectrophoretic trapping field is applied. This field centers the cell in the microcage, allowing for the measurement of one cell at a time. Each frequency results in a distinct rotation speed, depending on the cell and medium characteristics. Cell rotation is recorded optically for 3 seconds at a frame rate of 15 frames per second, totaling 1 minute and 3 seconds per cell. Graphs plotting the applied frequencies against the rotation speed generate electrorotation spectra. In the applied frequency range, the cell rotates in the opposite direction to the phase shift of the electric field, so the rotation speed is considered negative. The negative peak of a spectrum represents the maximum speed of rotation and is used to normalize the spectra between 0 and -1.

After a successful measurement, the cell is discarded by turning off the electric field and resuming the flow to capture the next cell. Throughout the experiment, cells remained suspended in a low conductivity medium of 200 mS/m and a controlled osmolarity of 300 mOsmol/kg.

Cell viability is monitored on-chip using a Live/Dead cytotoxicity and viability assay. The assay uses Calcein Acetoxymethyl ester (Calcein AM), which is converted by intracellular esterases to emit green fluorescence in viable cells, and ethidium homodimer-1 (EthD-1), which penetrates cells with compromised membranes and binds nucleic acids, producing a red fluorescent signal indicative of membrane damage. This method enabled us to distinguish between live and dead cells. Cells that are negative for Calcein AM and/or positive for EthD-1 are excluded from the datasets, ensuring that only viable cells are measured.

### Electrorotation results of membrane-compromising drugs mimicking the membrane damage by neurotoxic protein aggregates

Since no comparable experiments on the membrane capacitance of cells exposed to amyloidogenic proteins have been performed, it is still unknown what specific membrane capacitance changes to expect. Thus, initial experiments were designed to evaluate the system’s ability to quantitatively detect variations in the membrane when cells from the neuroblastoma line M17 were incubated with the membrane-damaging toxins Miltefosine (hexadecylphosphocholine) and Lovastatin, mimicking the membrane damage resulting from exposure to aSyn aggregate forms. Miltefosine is a synthetic phospholipid analogous to phosphatidylcholine, the major constituent of the cell membrane. It inhibits de novo membrane synthesis, thereby disrupting membrane integrity. Lovastatin, on the other hand, inhibits HMG-CoA reductase, the enzyme responsible for mevalonate production, leading to cell death.

When comparing our results to the membrane-compromising drugs, we see a similar trend in the relative shifts, both considering the spectra and the capacitances (Fig. 2). Membrane capacitances decrease significantly when cells are exposed to the drugs compared to the M17 reference measurement. Similarly, the shift in electrorotation spectra shows a clear shift to the right, resembling a decrease in membrane capacitance, which is also the case for EtHd-1 positive cells (dead cells). Another interesting comparison is between dead and alive cells (Fig. 2ab).

**Fig. 2.**
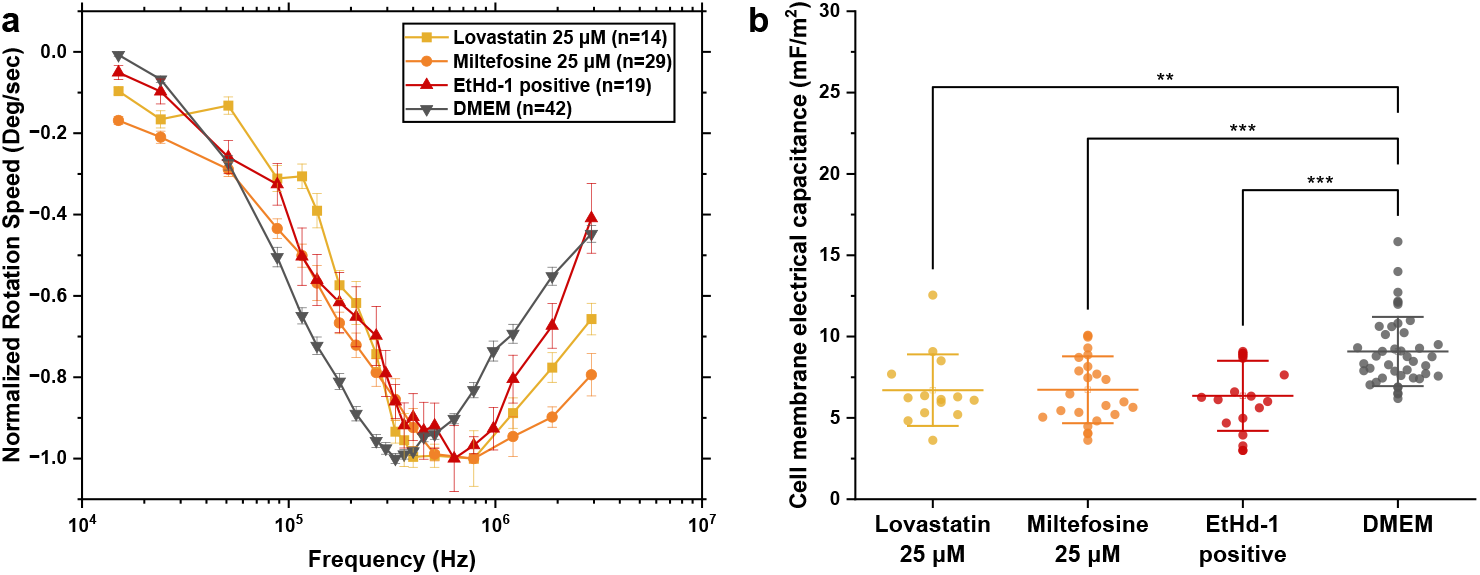
Electrorotation results of M17 cells treated with membrane-compromising toxins. **a** Averaged and normalized electrorotation spectra, with SEM as error bars, of M17 neuroblastoma cells in culture media as control (grey) or incubated with two different toxins (Miltefosine, orange, and Lovastatin, yellow) mimicking the membrane damage elicited by aSyn. The data in red represent dead cells positive to ethidium homodimer-1 (EthD-1). **b** Membrane capacitance values for individual cells. Points in scatter plots represent the membrane electrical capacitance of individual cells recorded in three independent experimental sessions (bars are means ± STDEV). A one-way ANOVA test (with Tukey test for paired comparisons) was performed among all the conditions (^*^ p≤0.05, ^**^ p≤0.01,^***^ p≤0.001).

### Validation of aSyn membrane association before electrorotation

Prior to introduction into the chip and measurement by electrorotation, Neuroblastoma M17 cells are incubated with monomeric, oligomeric and fibrillic forms of aSyn and a mixture of monomers and fibrils for timespans of 0, 3, and 6h. This was done to confirm the presence of the proteins on the cell membrane and ensure that we are measuring membrane toxicity related to aSyn interaction with the membrane and that all cells are exposed to the proteins at the chosen time points. We allow the cells to adhere by plating them 24h before incubation with the proteins. Fig. **3** shows the three different time points of incubation with a mixture of purified monomers and fibrils. We observe that the proteins are retained on the surface of all visible cells at 3h incubation and 6h incubation of all three different conditions, as confirmed by the imaging of the fluorescently labelled species (monomers with Atto647 and fibrils with Atto488). This indicates that the chosen concentrations are sufficient to ensure robust protein–cell membrane interactions,

**Fig. 3.**
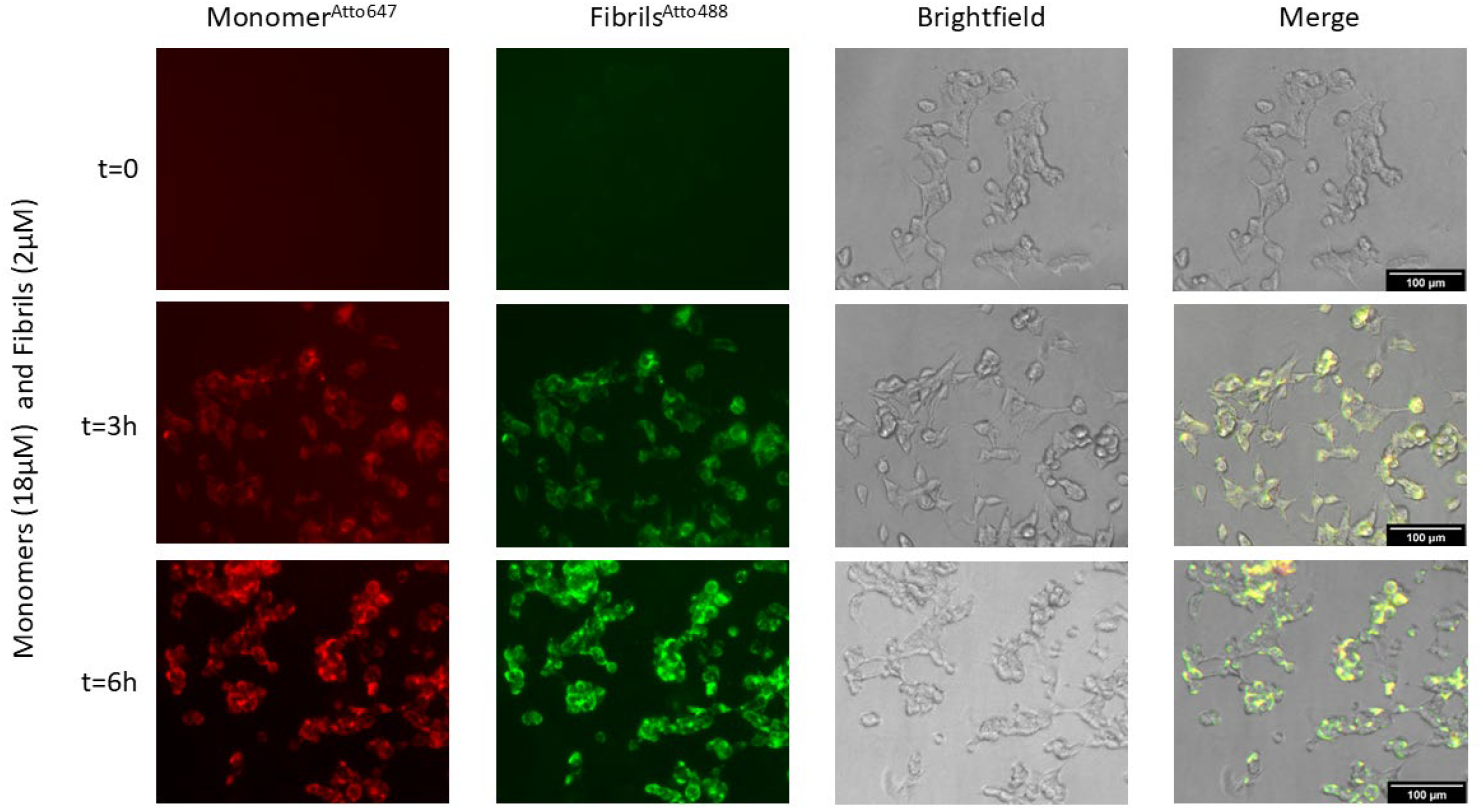
M17 neuroblastoma cells incubated with monomers and fibrils of α-Synuclein for 0h, 3h, and 6h. Monomers are labeled in Atto 647, and Fibrils in Atto 488. Cells were imaged in the TRITC fluorescence channel relative to Atto 647 (red), FITC for Atto 488 (green), and in brightfield; the panel on the right shows the merging of the three channels. The scale bar equals 100 µm.

As expected, the 0h condition did not show any observable signal as the species did not have time to interact with the cells yet. We used concentrations of monomers (18 µM) and fibrils (2 µM) similar to those previously reported by Mahul-Mellier *et al*.^10^ to induce toxic effects, as a mixture, after 48h *in vitro* using dye-exclusion assays. For the oligomeric forms, we used a concentration of 250 nM (based on unpublished data from the Lashuel lab).

### Verification of the presence of aSyn fibrils on the cell membrane during experimentation

The presence of aSyn fibrils on the cell membrane during rotation in one of the five electrorotation cages (Fig. 4ab) can be observed in Fig. **4**c for the fibrils-only condition and in Fig. **4**d for the monomers and fibrils mixture. Furthermore, no difference in Atto488 fluorescence intensity between the fibrils condition and the monomers and fibrils combination was determined. This indicates that the difference in membrane capacitance is not solely dependent on the presence and concentration of the protein at the membrane but rather their different interactions with the membrane.

**Fig. 4.**
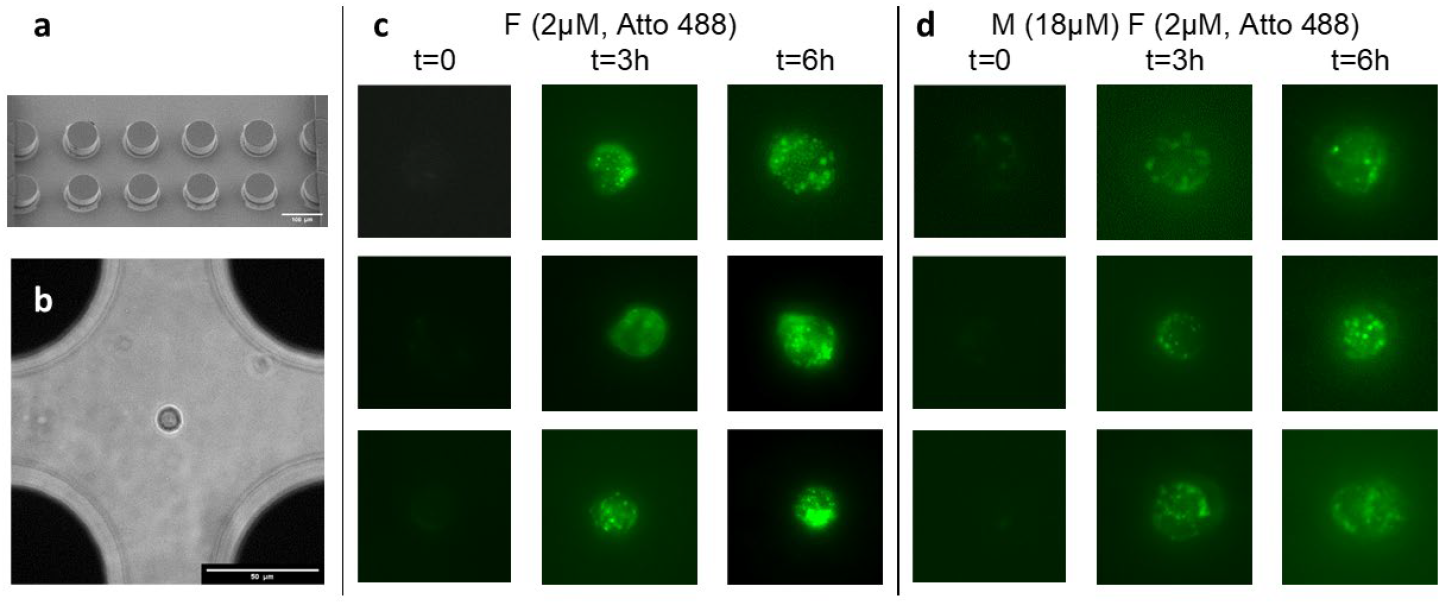
Verification of presence of fibrils inside the microfluidic chip. **a** Microcage array for parallel analysis of cells (Scale bar 100 µm). **b** Image of a neuroblastoma cell captured in a microcage. (Scale bar 50 µm). **c** M17 neuroblastoma observed in an electrorotation microcage after prior incubation with Atto 488-labeled fibrils (F) (2 µM) for 0, 3h, and 6h. **d** M17 neuroblastoma observed in an electrorotation microcage after prior incubation with monomers (M) (18 µM) and the Atto 488 labeled fibrils (2 µM) for 0, 3h, and 6h.

### Monomers and fibrils alone have no impact on membrane electrical capacitance

To assess by electrorotation the membrane damage caused by different forms of aSyn aggregation, we examined the effects of oligomers, monomers, fibrils, and a mixture of monomers and fibrils at various incubation times (0h, 3h, and 6h). Membrane damage was analyzed by evaluating shifts in the electrorotation spectra and changes in membrane capacitance compared to the electrokinetic response of a control condition. As a positive control for membrane damage, we used the membrane-compromising drugs Miltefosine at 25 µM, which inhibits de novo membrane synthesis, thereby mimicking membrane disruption and Lovastatin at concentrations of 25 µM, which alters the cell membrane indirectly by lowering cholesterol and isoprenoid levels, which affects membrane fluidity, lipid raft integrity, and membrane-associated signaling proteins.

The normalized electrorotation spectra and membrane capacitance values of the monomers (Fig. **5**a) revealed no measurable changes in the spectra of cells exposed to monomers at the three incubation time points compared to the negative control condition. The results were consistent across all three time points. Significant differences were only observed between the monomeric conditions and the positive control of cells treated with Miltefosine (Fig. **5**b).

**Fig. 5.**
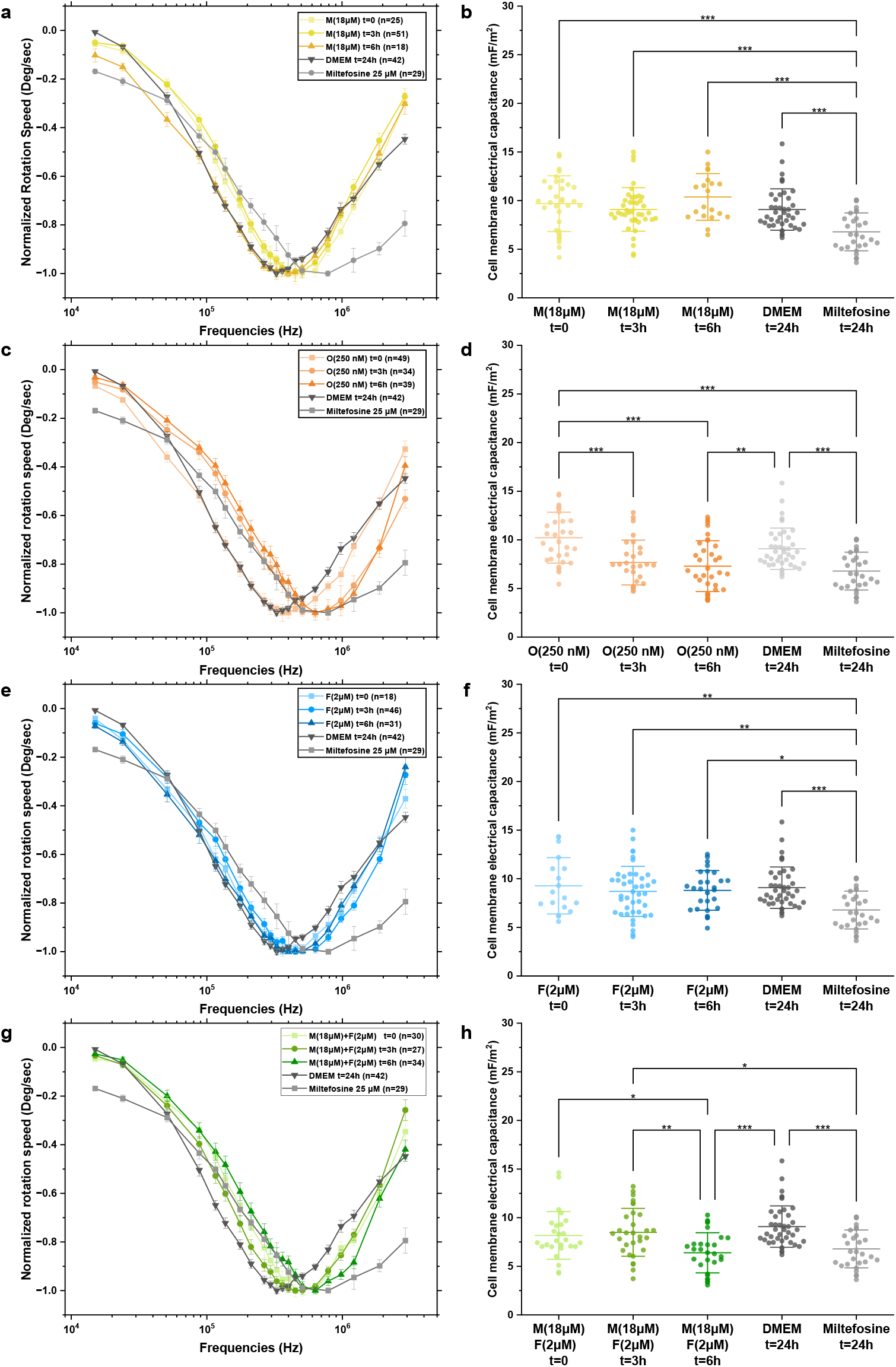
Electrorotation results for M17 cell incubated with different aSyn aggregate forms. **a**,**c**,**e**,**g** Averaged and normalized electrorotation spectra (bars are means ± S.E.M.) of M17 neuroblastoma cells incubated with the monomers (18 µM), oligomers (250 nM), fibrils (2 µM) and a combination of monomers (M, 18 µM) and fibrils (F, 2 µM) of human WT aSyn at three different time points (0, 3h and 6h) and their corresponding membrane capacitances extracted with the single-shell model **b, d, f, h**. In all graphs Miltefosine (25 µM) serves as a positive control and DMEM as a negative control. Points in scatter plots represent the membrane electrical capacitance of individual cells recorded in three independent experimental sessions (bars are means ± STDEV). A one-way ANOVA test (with Tukey test for paired comparisons) was performed among all the conditions (^*^ p≤0.05, ^**^ p≤0.01,^***^ p≤0.001).

Similarly, the electrokinetic responses of cells exposed to fibrils at different incubation times overlapped and matched the negative control condition (Fig. **5**e). Like the monomeric form of aSyn, this lack of spectral change indicates that fibrils do not alter membrane electrical capacitance, irrespective of the incubation duration (Fig. **5**f).

### Oligomers and a mixture of monomers and fibrils affect the cell membrane

Interestingly, observing the spectra of cells treated with oligomers at different time points suggests a different impact on the cellular membrane (Fig. **5**c). A noticeable shift in the normalized spectra is evident at both the 3h and 6h conditions compared to the reference, indicating a difference in membrane properties. The peak is located at a higher frequency than the negative control condition, correlating with a decrease in membrane electrical capacitance and, thus, a loss in membrane integrity (Fig. **5**d). Interestingly, we observed the occurrence of oligomer impact after 3h and 6h incubation as the spectra shifted towards higher frequencies. This shift toward higher frequencies was not observed after 0 hours of incubation, indicating that the oligomers do not have an immediate impact on the membrane.

Furthermore, an impact on membrane characteristics and electrorotation spectra was observed when treating neuronal cells with a mixture of monomers and fibrils (Fig. **5**g). The shift in the monomer and fibril mixture condition appears only after 6h of incubation, as a shift compared to the negative control and previous time points. On the other hand, no significant difference is seen between 0h, 3h, and the negative control. For ease of comparison, we plotted the same data shown previously (Fig. **5**), considering the results at 6h incubation for all measured conditions and controls. Comparing the spectra of all aggregate forms after 6h shows an overlapping shift for the oligomer condition and the monomers and fibrils combination (Fig. 6a). The overlapping spectra align with the significant decreases in membrane capacitance observed in these conditions, as well as with the positive Miltefosine control (Fig. 6b). Notably, there is a similar decrease in membrane capacitance for the membrane-compromising drug Miltefosine and the aSyn toxic conditions (oligomers and monomers, and fibrils).

**Fig. 6.**
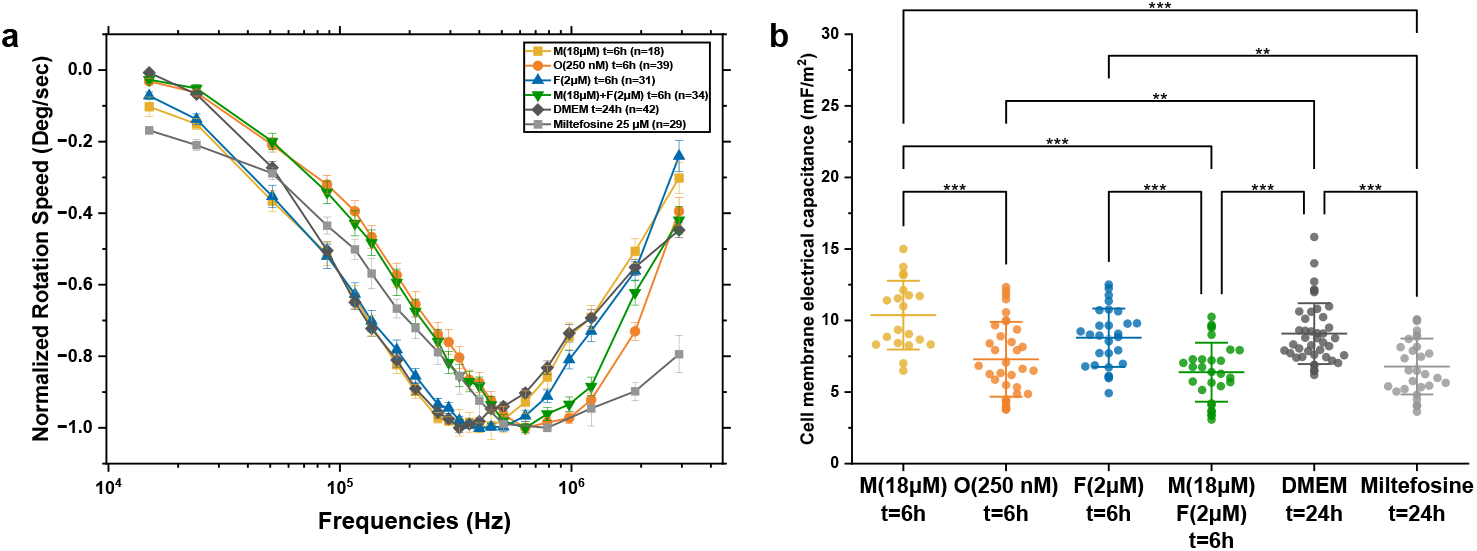
Electrorotation results for M17 cell incubated with different aSyn aggregate forms for 6 hours. **a** Averaged and normalized electrorotation spectra (bars are means ± S.E.M.) of M17 neuroblastoma cells incubated with different aggregate forms of aSyn after 6h incubation and their corresponding membrane capacitances **b**. Points in scatter plots represent the membrane electrical capacitance of individual cells recorded in three independent experimental sessions (bars are means ± STDEV). In both graphs Miltefosine (25 µM) serves as a positive control and DMEM as a negative control. A one-way ANOVA test (with Tukey test for paired comparisons) was performed in between all the conditions (^*^ p≤0.05, ^**^ p≤0.01,^***^ p≤0.001).

The 6h-fibril and monomer mixture incubation conditions lack this shift and overlap with the untreated negative control, indicating these forms are non-toxic. Furthermore, they show a significant increase in membrane capacitance when compared with the positive control and no significant difference with respect to the negative control.

## Discussion

This study demonstrates the power of microfluidic electrorotation as a sensitive and label-free method for assessing membrane integrity at the single-cell level. While our investigation centered on aSyn, a key protein in PD pathology, the broader significance lies in the application of electrorotation to quantitatively study membrane-related toxicity across a range of pathological conditions.

We successfully leveraged this platform to differentiate the effects of various aSyn aggregate forms (monomers, oligomers, fibrils, and a mixture of monomers and fibrils) on cellular membranes. Importantly, electrorotation allowed us to capture early timepoint resolved changes in membrane electrical capacitance, which is a direct indicator of membrane structure and function. We were able to observe toxicity onset as early as 3h, and to detect differences in the amount of damage caused by different aSyn forms.

Our findings demonstrate that neither monomers nor fibrils alone induce membrane-related toxicity up to 6h of incubation. In contrast, purified oligomers and a mixture of fibrils with monomers significantly reduced membrane electrical capacitance, similar to the effects observed with membrane-disrupting drugs like Miltefosine and Lovastatin. These results support existing literature identifying oligomers as the most toxic aSyn species due to their pore-forming^28^ and detergent-like properties^33^. While fibril toxicity remains controversial^30,35,36^, our data support the view that fibrils alone are non-toxic and require the presence of monomers to exert membrane-disruptive effects^10^. This is likely due to monomer-facilitated fibrillization at the membrane, consistent with prior studies^10^. The observed decrease in capacitance and the spectral peak shift point to compromised membrane integrity as the underlying mechanism. Additionally, the time-dependent nature of toxicity, emerging after 3h for oligomers and 6h for fibril-monomer mixtures, may reflect differences in their modes of action, such as oligomer membrane insertion versus fibril growth-mediated membrane disruption. Further investigation is needed to elucidate the concentration-dependent effects and distinguish between pore formation and detergent-like disruption.

Unlike traditional bulk assays, our single-cell microfluidic design enables the analysis of dielectric properties under well-controlled conditions. The incorporation of a Galton array with electrorotation microcages facilitates measurements across multiple cells per experiment and in parallel, improving statistical power while preserving single-cell detail. This precision is crucial for detecting heterogeneous responses that may otherwise be masked in population-averaged measurements.

Beyond aSyn, the approach and findings reported here have implications for other pathological proteins implicated in Alzheimer’s disease (e.g., Aβ peptides)^3^ and Huntington’s disease (e.g., mutant huntingtin)^38^, and are known to interact with lipid membranes, potentially compromising their integrity. Electrorotation could be used to systematically compare the membrane-disruptive capacity of these aggregates and to screen for therapeutic agents that restore membrane stability. Moreover, the method is broadly applicable to the study of membrane-targeting agents such as bacterial toxins^39^, chemotherapeutic drugs^40^, or environmental pollutants^41^. For example, electrorotation could detect pore-forming activities of bacterial virulence factors or monitor early-stage damage from detergent-like compounds without relying on invasive fluorescent labels or endpoint assays.

Although we cannot yet distinguish between specific mechanisms (e.g., pore formation vs. detergent-like effects), this method robustly identifies the functional outcome, a compromised membrane capacitance. This enables the rapid screening of pathological agents for membrane damage, as well as the identification of inhibitors that block toxin- or protein aggregate-induced membrane damage, which could serve as potential therapies for the treatment of PD and other amyloid-related disorders.

In summary, our results validate electrorotation as a powerful biophysical platform for probing membrane dynamics in real time. The ability to detect subtle, time-dependent changes in electrophysiological properties provides a new lens for studying early toxicity mechanisms, opening avenues for diagnostics, drug development, and fundamental membrane biology. The technology’s adaptability makes it a valuable asset for a wide range of biomedical research, far beyond its demonstrated utility in aSyn-related neurodegeneration.

## Materials and methods

### Microfabrication

Detailed microfabrication was previously described by Kilchenmann *et al*.^42^. The fabrication process involved sputtering a Ti/Pt/Ti (20/200/20 nm) layer onto a planar wafer using a Pfeiffer Spider 600 system. Subsequently, ion beam etching (Veeco Nexus IBE350) was employed to pattern connection lines. A SiO_2_ layer (300 nm) was sputtered to insulate the wires, and vias were etched into the insulation layer using the SPTS APS Dielectric Etcher. To create the 3D electrodes, a 50 µm SU-8 (MicroChem 3025) scaffold was patterned on the exposed metal pads, which was then covered with a Ti/Pt (20 nm/200 nm) metal layer by sputtering. Ion beam etching without an etching mask at an angle of 0° perpendicular to the wafer surface removes all the metal on the wafer except on the side walls of the electrodes. A second layer of 50 µm SU-8 was coated and patterned on the wafer to form the wide microfluidic channel, support structures, and the Galton array. The chip assembly was completed by irreversibly bonding PDMS to the SU-8 channel using APTES^43^. An SEM image of the finalized chip and a device schematic are presented in Fig. **1**.

### Protein expression

Recombinant human wild-type aSyn was expressed in E. coli using pT7-7 plasmids. aSyn was then purified by anion-exchange chromatography followed by reverse-phase high-performance liquid chromatography (HPLC). Lyophilized aSyn monomers were dissolved in 50 mM Tris and 150 mM NaCl, and the pH was adjusted to 7.5 with 1 M sodium hydroxide. The solution was then filtered using 100 kDa molecular weight cut-off filters. The purity and identity of the recombinant proteins were confirmed as described by Fauvet et al.^44^ and the concentrations where determined using a Pierce™ BCA Protein Assay (Thermo Fisher Scientific). Detailed characterization of all protein material used in this study can be found in recent publications from our group using the same protein material^10,45,46^.

Fluorescent labelling of aSyn monomers used for figure 3 was carried out as previously described^10^. Recombinant V16C-α-synuclein was dissolved in 200 mM Tris buffer, pH 7.0 (Trizma hydrochloride solution, Sigma-Aldrich). To reduce disulfide bonds, two molar equivalents (final concentration 200 μM) of TCEP (Sigma-Aldrich) were added, and the solution was incubated at room temperature (RT) for 5 minutes. Subsequently, Atto 647 maleimide (Atto-TEC) was added to the protein solution at a final concentration of 500 μM. The labeling reaction was monitored by electrospray ionization mass spectrometry (ESI-MS), and after 1 hour, >95% of V16C-α-synuclein was successfully conjugated with the maleimide dye. Unreacted dye was removed by gel filtration using a PD-10 column (Fisher Scientific AG, Lucens, Switzerland) according to the manufacturer’s protocol. Briefly, the column was equilibrated with 50 mL of deionized water, and the protein solution (diluted to 2.5 mL) was applied and eluted with water. Eluted fractions containing labeled α-synuclein were identified by SDS-PAGE, pooled, and lyophilized. The correct molecular mass of the labeled protein was confirmed by MALDI-TOF-MS and ESI-MS. Labeling efficiency and purity were further verified by SDS-PAGE and UV-vis spectroscopy.

Unmodified WT aSyn oligomers were prepared as previously described^47,48^. For oligomer preparation, after the HiPrep column, the aSyn-positive fractions were dialyzed against de-ionized water at 4°C overnight to remove salts, subsequently snap-frozen, and lyophilized. Subsequently, 60 mg of the lyophilized protein was dissolved in 5 mL of PBS buffer (pH 7.4) and incubated at 900 rpm on an orbital shaker at 37°C for 5h. Later, the sample was centrifuged at 12,000 g at 4°C for 10 min to remove any insoluble species. The supernatant was then loaded onto a HiLoad 26/600 Superdex 200 preparation grade (GE Lifesciences) column pre-equilibrated with PBS buffer (pH 7.4). Protein was eluted in 2.5-ml fractions at a flow rate of 1 ml/min. All fractions were further analyzed by Sodium Dodecyl Sulfate-Polyacrylamide Gel Electrophoresis (SDS-PAGE) and transmission electron microscopy (TEM), and the oligomer-positive fractions are snap-frozen and stored at -20°C until needed for experiments or further analysis.

WT aSyn fibrils were similarly prepared as previously stated^47–49^. For the preparation of fibrils, 4 mg of aSyn monomers was diluted in 600 μl of PBS (pH 7.4), and the pH was adjusted to 7.4. The monomeric aSyn solution was filtered through 0.2-μm filters (Merck, SLGP033RS) using ultracentrifugation (12,000 g, 10 min, 4°C) before being incubated on an orbital shaker (1000 rpm) at 37°C for 5 days. The fibril formation was evaluated by SDS-PAGE and Coomassie staining before the sample was sonicated on ice with a fine tip (Sonics Vibra cell) for 20 s, at 20% amplitude, a pulse of 1 s ON/1 s OFF (Sonic Vibra-Cell, Blanc-Labo, Switzerland). The number of monomers and oligomers released from sonicated fibrils was quantified using the filtration protocol that we developed^47^. Sonicated aSyn fibrils were characterized by TEM and snap frozen. They can be stored at -80°C until needed.

Fluorescent labelling of aSyn fibrils used for figures 3 and 4 was carried out following as previously described^53^. Fibrils were diluted to 250 µM in 500 µL PBS (pH 7.5). An equimolar amount of Atto 488 maleimide (Atto-Tec, Switzerland) was added, and the mixture was incubated overnight at 4 °C. Labelled PFFs were pelleted by ultracentrifugation at 100,000 g for 1h at 4 °C. The supernatant was removed, and the pellet was resuspended in PBS; this washing step was repeated until excess dye was eliminated. Successful labelling was confirmed by SDS–PAGE and fluorescence scanning (Typhoon FLA 7000, GE Healthcare; excitation/emission: 400/505 nm). The labelled fibrils were sonicated (four cycles of 5 s at 20% amplitude; Sonic Vibra Cell, Blanc Labo, Switzerland). Aliquots were snap-frozen in liquid nitrogen and stored at – 80 °C. Fibril structure and biophysical characteristics were verified by TEM, SDS– PAGE with Coomassie staining, and thioflavin T assays.

### Cell preparation and treatment

BE(2)-M17 human neuroblastoma cells were cultured in complete Dulbecco’s Modified Eagle’s Medium (DMEM (Gibco, Thermo Fisher Scientific) supplemented in 10% Fetal Bovine Serum (Gibco, Thermo Fisher Scientific) and 1% of an antibiotic-antimitotic solution (L-Glutamine-penicillin-streptomycin, Thermo Fisher Scientific). 24h before the experiments, cells were plated in 6-well plates at a concentration of 200’000 cells in 1.5 mL of complete DMEM. aSyn proteins were added for different time points at concentrations of 2 µM for the fibrils, 18 µM for the monomers, and 250 nM for the oligomers. Similarly, Miltefosin (Sigma Aldrich) and Lovastatin (Sigma Aldrich) where added at concentrations of 25 µM for 24h. At the indicated time, cells were detached using 1X Trypsin (Thermo Fisher Scientific). After centrifugation at 1200 rpm and supernatant removal, all cells were resuspended in low conductivity buffer (8.6% dextrose and 0.3% sucrose) with an adjusted conductivity of 200 mS/m using 1X PBS (Gibco, Thermo Fisher Scientific). Cell concentration was maintained at approximately 200,000 cells per mL. Cell viability was monitored on-chip using kit containing Calcein AM and Ethidium homodimer-1 (LIVE/DEAD™ Viability/Cytotoxicity Kit, for mammalian cells, Thermo Fisher Scientific). Prior to experiments, the microchips were primed by flushing with 2 mL of the 200 mS/m conductivity medium with 1% Bovine Serum Albumin (Thermo Fisher Scientific) to avoid cell adherence to the channel walls.

### Measurement procedure

Cells were injected into the device using a pressure pump at pressures ranging from 1 to 10 mbar. Applying an alternating current (100 kHz) electric voltage signal created dielectrophoretic barriers at the entrance and exit of the array, trapping cells within the microcages. After trapping, the cells were subjected to a rotating electric field (3 V amplitude, swept frequency, and 90° phase shift between neighboring electrodes) for electrorotation. The rotation speed was recorded through microscope imaging, and a rotating electric field was swept at 21 different frequencies between 15 kHz and 2.14 MHz. Chips can be reused up to 10 times if cleaned by flushing consecutively 1 mL PBS, 1% sodium hypochlorite through the chip, and DI water to remove biological material.

### Dielectrophoretic force and electrorotation phenomenon

The dielectrophoretic force *F*_*DEP*_, stabilized the cell in the center of the microcage by balancing the drag force of the fluid flow. It is expressed as ^50^:

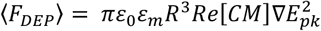

where *ε*_0_ is the absolute permittivity and *ε*_*m*_ the permittivity of the surrounding medium. R is the Radius, and 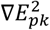 is the gradient of the electric field amplitude. Re[CM] is the real part of the Clausius-Mossotti factor and describes the dielectric properties of the cell and the surrounding media.

Rotation of the cell is induced by a consecutive phase shift of 90° for each respective electrode. The speed of electrokinetically-induced rotation is given by^50^:

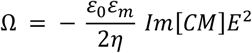

*Im*[*CM*] depends on the model used for the cell, which is, in our case, the single shell model, and is given by^21^:

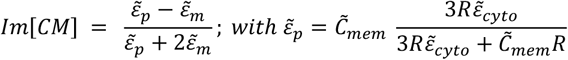

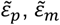 and 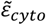 are the complex permittivities of the particle, suspending medium, and cytoplasm, respectively. Complex permittivities are given as *ε* ∗= *ε* − *iσ*/*ω*, and the complex capacitance 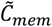 defined as 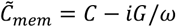 with G being the membrane conductance.

### Data analysis, extraction of membrane capacitance, and data representation

To get sufficient data points for fitting the electrorotation spectra to the model above, we apply a set of 21 different frequencies and a voltage of 3 V_*p-p*_ while imaging the cells at a frame rate of 15 frames per second during 3 seconds per frequency. In the next step, the cell rotation and radius are extracted using an in-house LabVIEW program. It identifies the speed of rotation using a pattern-matching algorithm and extracts the radius of the measured cells.This results in a rotational spectrum acquired for each cell measured. From there, we use a custom MATLAB script to extract the membrane’s electrical capacitance. For computation, we use the nonlinear least-squares solving function *lsqcurvefit* in MATLAB.

Finally, present our data by representation of both the membrane capacitance acquired for each cell, but also an averaged and normalized electrorotation spectra to be able to directly compare the rotational behavior of different conditions without the impact of any additional parameters other than the rotation of the cell. Typically, individual cells have different rotation speeds. Thus, the comparison of raw data would be chaotic with simple averaging of rotation speeds. Due to this, we normalized each of the acquired spectra at the minimum speed of rotation (normalization to -1). Finally, we normalize a second time to keep the negative peak of the spectra at -1. Comparing the negative peak of different conditions gives us valuable insight into the capacitive behavior of the membrane, as a general shift of the minimum peak toward higher frequencies equals a decrease in membrane capacitance, and toward lower frequencies equals a reduction in membrane capacitance.

## Acknowledgements

This research was funded by the Swiss National Science Foundation (205321 179086).

## Author contributions

T.R. designed and fabricated the device, designed and performed experiments, and wrote the manuscript. A.K. contributed to device design., N.M and F.R.E. helped with data acquisition and analysis processing; A.L.M.M helped with experimental design and proofreading. H.L and C.G. supervised the work, acquired funding, and reviewed the draft of the manuscript.

